# Modelling the movements of organisms by stochastic theory in a comoving frame

**DOI:** 10.64898/2026.02.11.705365

**Authors:** Norberto Lucero-Azuara, Rainer Klages

## Abstract

Imagine you walk in a plane. You move by making a step of a certain length per time interval in a chosen direction. Repeating this process by randomly sampling step length and turning angle defines a two-dimensional random walk in what we call *comoving frame* coordinates. This is precisely how Ross and Pearson proposed to model the movements of organisms more than a century ago. Decades later their concept was generalised by including persistence leading to a correlated random walk, which became a popular model in Movement Ecology. In contrast, Langevin equations describing cell migration and used in active matter theory are typically formulated by position and velocity in a fixed Cartesian frame. In this article, we explore the transformation of stochastic Langevin dynamics from the Cartesian into the comoving frame. We show that the Ornstein-Uhlenbeck process for the Cartesian velocity of a walker can be transformed exactly into a stochastic process that is defined *self-consistently* in the comoving frame, thereby profoundly generalising correlated random walk models. This approach yields a general conceptual framework how to transform stochastic processes from the Cartesian into the comoving frame. Our theory paves the way to derive, invent and explore novel stochastic processes in the comoving frame for modelling the movements of organisms. It can also be applied to design novel stochastic dynamics for autonomously moving robots and drones.

## I. INTRODUCTION

Organisms living at very different spatio-temporal scales, from moving in the microworld to foraging across the surface of the earth, display highly complex, random-looking migration patterns [1]. By now there exists a wealth of experimental recordings of these patterns for a huge variety of species, from insects [2–4] to fishes [5–7], birds [8–10], mammals [11–13] and even humans [14–16]. Novel biologging techniques developed over the past decade are delivering bigger and more precise data sets [17–19]. These developments pose the fundamental challenge to understand the experimentally recorded organismic movement patterns by constructing mathematical models from data [20],

Around 1905 Ross and Pearson introduced random walks for modelling the migration of organisms, described in terms of step length and turning angle with respect to the previous step for movements in a plane [21–23]. In the simplest case, one may choose a constant step length while the turning angle is sampled independently and identically distributed from a uniform probability distribution, reflecting the randomness of organismic movement [23, 24]. This simple model became popular as the drunkard’s walk [23, 25], since its dynamics does not contain any memory, which mathematically corresponds to a Markov process [26, 27]. It is at the heart of describing diffusive spreading in nature, technology and society [28]. For decades simple random walks have been used to model organismic movements [29] until in the 1980’s they were generalised by sampling the turning angle from a unimodal distribution, which imposes a correlation between two subsequent steps modelling one-step persistence in organismic motion [2]. In addition, one may choose the step length as an independent and identically distributed random variable. This model became known as a correlated random walk (CRW) [30]. Together with state space, [31, 32], hidden Markov [31, 33] and other stochastic models [30, 34] CRWs form the theoretical backbone of *Movement Ecology* (ME), a field founded in 2008 [35, 36] mainly driven by experimental biologists [17, 18], which endeavours to understand the movements of organisms, especially on larger scales, in view of their interactions with the environment. One may ask, however, whether including one-step persistence is sufficient to fully understand the movements of organisms, which often feature long-term memory way beyond a single step.

Another fundamental approach to model movement patterns draws on the observation that they may look similar to the Brownian motion of a tracer particle in a fluid, described by the famous Langevin equation (LE) [37]. This equation is based on Newton’s second law by decomposing the force acting onto a Brownian particle into Stokes friction and random collisions with the surrounding molecules, modelled by Gaussian white noise. Formulated in terms of Cartesian position and velocity of a moving particle, it is sometimes called Newton’s law of stochastic physics. Mathematically, the LE represents a Markovian Ornstein-Uhlenbeck (OU) process [38] for the velocities of the Brownian particle [39]. Like simple random walks, OU processes have been, and still are, widely used to model organismic movement [31, 40]. In parallel, however, Langevin dynamics was generalised to reproduce the non-trivial dynamics of migrating cells [41–45] and active particles [46–49].

Crucially, CRWs and the (generalised) LEs referred to above are formulated in very different coordinate frames. What we call *comoving frame* coordinates traces back to Ross and Pearson’s original formulation of two-dimensional random walks, for which they used step length and turning angle, as illustrated in Fig. 1. Note that in this figure the step length is replaced by the speed of an organism, where the speed is defined as step length per time interval. In that sense, with comoving frame we denote a frame attached to the center of mass of an organism cotranslating and -rotating with its movements, whose abscissa is aligned with the velocity of an organism, and the rotations of this velocity vector yield the associated turning angle. Comoving frame coordinates are biologically very well motivated, as it seems natural to think of higher-dimensional movements of organisms in terms of step length and turning angle [21, 23, 24]. Correspondingly, step length distributions are intimately related to step selection functions and associated concepts widely applied in ecology and conservation to experimentally characterize the movements of organisms [50–52]. LEs, in contrast, are typically defined in terms of Cartesian position and velocity of a moving particle [25–27, 37]. This description makes perfect sense for representing the passive motion of a tracer particle driven by collisions from the surrounding molecules in a fluid that as a whole is at rest in a fixed Cartesian frame. It also simplifies the theoretical analysis of these equations, especially if they are generalised by including memory kernels [41–45]. However, one may fundamentally question whether the Cartesian approach is correct for modelling self-propelled, active movement that is generated intrinsically by an organism itself, instead of having a passive particle solely driven extrinsically by interactions with the environment [1]. One may argue that active fluctuations emerging internally within an organism should rather be modelled by a stochastic process defined *self-consistently* in a frame comoving with this organism, i.e., without explicitly involving any other coordinates than speed and turning angle, and not by noise or friction terms somewhat formulated externally with respect to a fixed Cartesian frame. This problem is indeed taken into account, to some extent, in some active particle models [47, 48, 53, 54], where upon closer scrutiny one popular type of them [48, 55] turns out to be identical to the CRW of ME [1]. However, as we will demonstrate in this article, to fully solve this problem one has to go one significant step further. There are thus many reasons to investigate merging the approaches of CRWs in a comoving frame and LEs in a fixed Cartesian frame for constructing a more general organismic movement model that combines advantages from both theories.

**FIG. 1.**
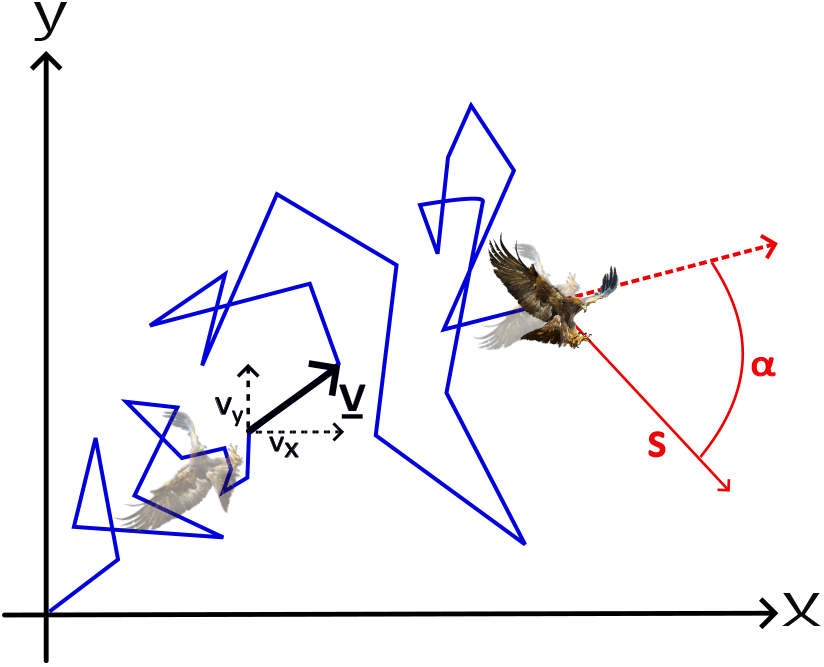
Time-discrete trajectory (blue) of an organism (the pictures show an eagle) in a fixed Cartesian frame (black horizontal and vertical lines). Also shown is the velocity of the eagle at some point along the trajectory in Cartesian coordinates *v*_*x*_, *v*_*y*_. In contrast, at the final point of the trajectory we represent the movement of the eagle in comoving frame coordinates (red) in terms of speed *s* and turning angle *α*, where the latter is defined as the angle between subsequent velocity directions (dashed, respectively bold red lines).

For constructing a stochastic model of bumblebee flights from experimental data, in 2013 Lenz et al. addressed this problem by fusing *ad hoc* generalised Les with CRWs in terms of two coupled stochastic differential equations for speed and turning angle, both of Langevin type but with *per se* arbitrary friction and noise terms [3]. All these terms were extracted from experimental data. Interestingly, the turning angle distribution was found to be unimodal, as in a CRW, all noise terms were correlated, and the relevant friction term was speed-dependent, as for a specific active particle model [46, 47]. Very recently, the same approach of formulating generalised Langevin dynamics in a comoving frame was exploited for experimentally steering superparamagnetic colloidal microrobots with tailored statistics generating non-Brownian anomalous diffusion, like fractional Brownian motion, by fine-tuning magnetic fields [56]. The underlying general framework thus promises to cross-link [1] the different big fields of ME [20, 35, 36], Active Particles [46–49] and Anomalous Diffusion [39, 57–60] for modelling and understanding the movements of organisms. This motivates to explore how to analytically derive the equations stipulated in [3] from first principles.

Our article solves this problem for the paradigmatic example of the OU process defined in the Cartesian frame. In Sec. II we first show that Cartesian and comoving coordinates are cross-linked by polar coordinates. We thus have to carefully distinguish between three different frames for our transformation of stochastic processes, which are the Cartesian, the polar and the comoving frame. As a tutorial warm-up exercise, in Sec. III we illustrate the transformation between these three different frames for the example of a simple Markovian random walk defined in the Cartesian frame by deriving corresponding stochastic models in the other two frames. Analytical results for all three models are compared with computer simulations. In Sec. IV we apply the same conceptual framework to analytically construct different models in the polar and in the comoving frame for the OU process defined in the Cartesian frame. Again all results are verified by computer simulations. We conclude with a summary of our main results, a wider embedding and an outlook to further research in Sec. V.

## II. STOCHASTIC PROCESSES IN TWO DIMENSIONS

In this section we establish the basic analytical framework for transforming two-dimensional stochastic processes from the Cartesian into the comoving frame by introducing three distinct frames of reference: the *Cartesian*, the *polar* and the *comoving* frame. Figure 1 shows a cartoon of an organism moving along a time-discrete random trajectory in a plane. In the fixed Cartesian frame, its velocity vector **v** is described by the two components (*v*_*x*_, *v*_*y*_), which specify the direction and magnitude of motion relative to a stationary reference system. By contrast, the comoving frame characterizes the same motion in terms of the speed *s* and the turning angle *α*, emphasizing changes in orientation relative to the direction of motion.

Figure 2 depicts the geometric relationships between these three reference frames by illustrating their different representations. A clear understanding of the transformations between Cartesian, polar, and comoving descriptions is essential for our subsequent systematic analysis of stochastic dynamics in two dimensions, and for interpreting the underlying physical and biological processes across these different modeling contexts. Extending the concept of stochastic processes evolving randomly over time from one to two dimensions introduces greater complexity and richness, due to considering a new degree of freedom. Mathematically, within our context a time-discrete two-dimensional stochastic process is typically represented by a vector **x**_*n*_ for the particle’s position coordinates at discrete time *n* ∈ ℕ_0_ in the Cartesian frame, characterized by probabilistic properties such as distributions, mean, variance, and correlations in both components *x*_*n*_ and *y*_*n*_. We now introduce these three reference frames in detail and define the different transformations between them.

**FIG. 2.**
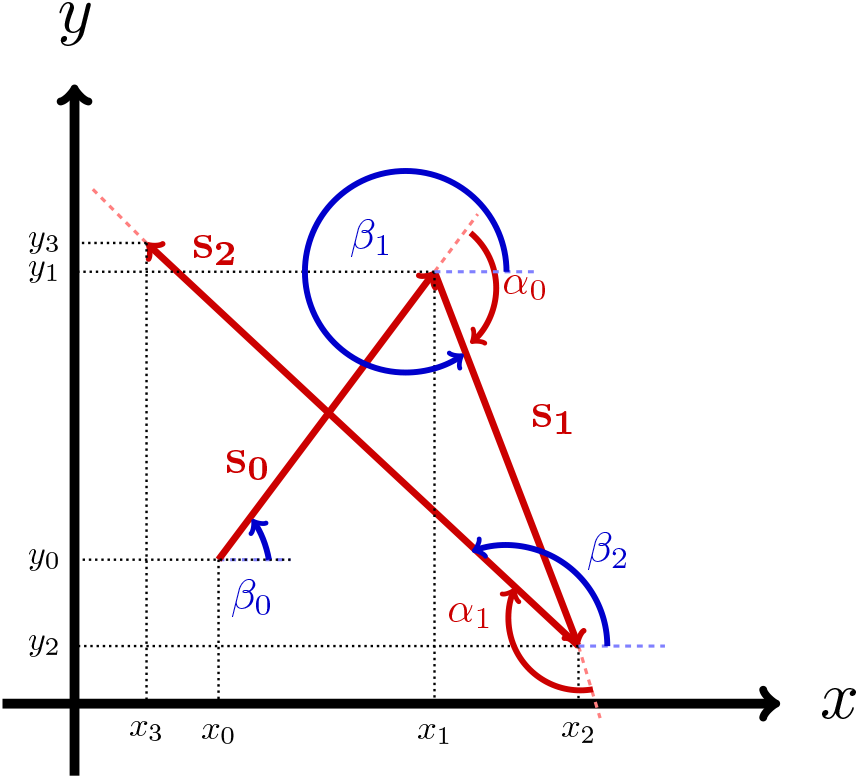
Conceptual illustration of the interplay between three different reference frames for a process over four discrete time steps *n* = 0, 1, 2, 3: Cartesian frame described by positions *x*_*n*_ and *y*_*n*_ (black), polar frame described by the orientation angle *β*_*n*_ (blue) and the speed *s*_*n*_ (red), and comoving frame described by the turning angle *α*_*n*_ and speed *s*_*n*_ (red).

### A. Frames of reference and transformations

#### 1. Cartesian Frame

This fundamental reference frame uses position coordinates (*x*_*n*_, *y*_*n*_) and velocity components (*v*_*x,n*_, *v*_*y,n*_) for describing a particle’s motion. Here and in the following we consider this frame to be at rest. For stochastic processes it is typically assumed that motion along the *x*-axis does not influence motion along the *y*-axis, and vice-versa, allowing for separate treatment of dynamics in each dimension. This simplifies the analysis for systems in terms of decoupled motion in different directions.

#### 2. Polar Frame

This frame is based on a polar coordinate transformation of the particle’s velocities, which is particularly useful for systems where radial and angular motion are key. Cartesian coordinates (*v*_*x,n*_, *v*_*y,n*_) are transformed into polar coordinates (*s*_*n*_, *β*_*n*_), where *s*_*n*_ is the absolute value of the vector velocity and *β*_*n*_ is the polar angle of this vector. The transformation is trivially given by

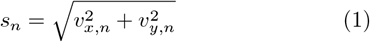

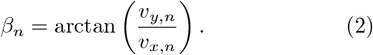

Conversely, to convert back to Cartesian coordinates we have

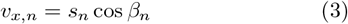

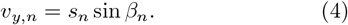

#### 3. Comoving Frame

This dynamic, particlecentered frame co-moves and -rotates with the particle, characterized by the particle’s *speed s*_*n*_ (as in Eq.(2)) and *turning angle α*_*n*_. It is defined with respect to the change of the time-discrete orientation angle of the vector velocities as

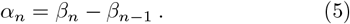

Here, *α*_*n*_ yields the turning angle between consecutive time steps *n* and *n* − 1. The inverse transformations from the comoving frame back to Cartesian velocity components are

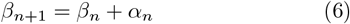

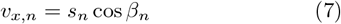

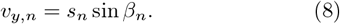

These equations reconstruct *v*_*x,n*_ and *v*_*y,n*_ from *s*_*n*_ and *α*_*n*_. The comoving frame is ideal for self-consistently describing intrinsic stochastic fluctuations driving a self-propelled particle, such as in active matter systems or organismic movement models, as we will show in the following.

The interplay between these three different frames of reference in terms of the respective transformations between them is summarised in Fig. 3.

**FIG. 3.**
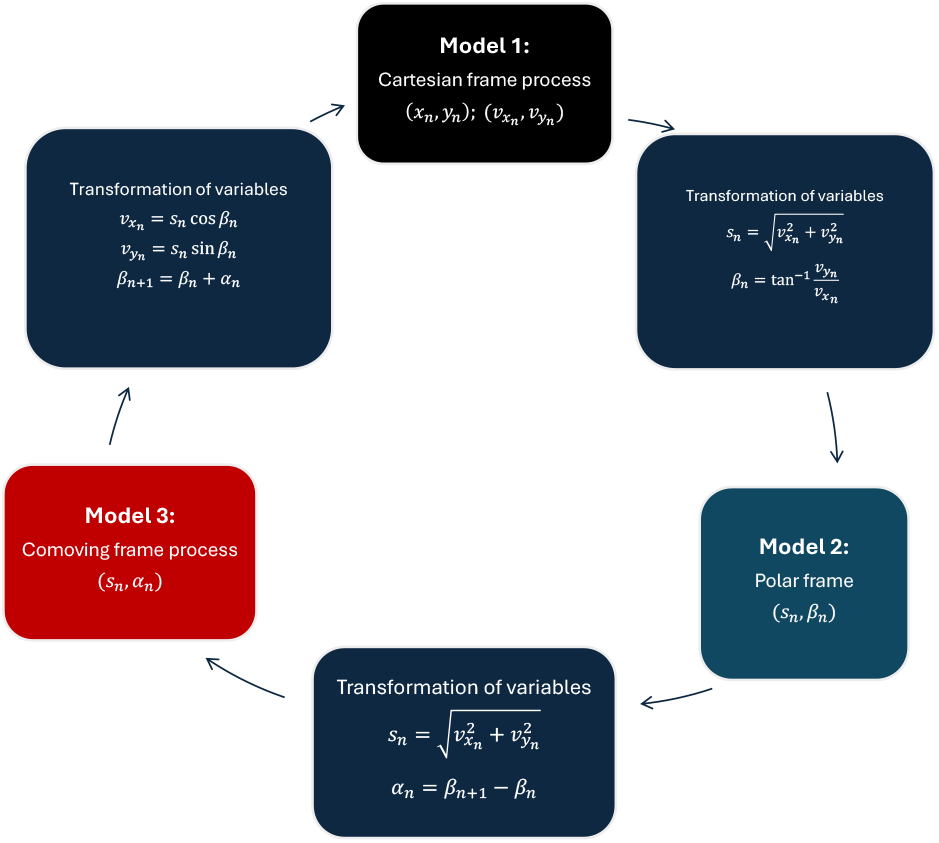
Full transformation rules between the three different frames of reference and the respective inverse transfromation.

## III. THREE MODELS FOR A SIMPLE RANDOM WALK IN TWO DIMENSIONS

In order to outline the basic principle of transforming a stochastic process between these three different frames of reference, in the following we first consider the example of a simple two-dimensional random walk. We start by defining the random walk in Cartesian coordinates. We then subsequently transform this process into polar and then into comoving coordinates, hence arriving at three different representations, or models, of this process in three different frames. The analytical results are compared with computer simulations. This analysis will pave the way to transform the OU process, which we do in Sec. IV.

### A. Model 1: Random Walk in the Cartesian frame

We first consider a two-dimensional time-discrete random walk defined by

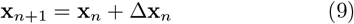

with increments Δ**x**_*n*_ = Δ*t***v**_*n*_, where the discrete velocities **v**_*n*_ = (*v*_*x,n*_, *v*_*y,n*_) are drawn from a Gaussian distribution *P*(*v*_*x*_, *v*_*y*_) with zero mean and variance *σ*^2^. This corresponds to an isotropic random walk in the plane, with independent and identically distributed (i.i.d.) steps. The Cartesian formulation thus describes the random walk in terms of Gaussian increments in each coordinate direction.

### B. Model 2: Random Walk in the polar frame

For transforming to polar coordinates, we use the speed *s*_*n*_ and orientation angle *β*_*n*_ defined by Eq.(2). In order to formulate a two-dimensional random walk in these coordinates, we need to transform the Cartesian Gaussian velocity distribution into polar coordinates. This change of variables is accomplished via using conservation of probability and the Jacobian determinant yielding

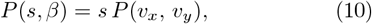

which in the isotropic Gaussian case of a normal distribution, *v*_*x*_, *v*_*y*_ ∼ 𝒩 (0, *σ*^2^), becomes

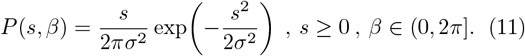

This factorizes into a Rayleigh distribution for the speed,

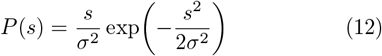

and a uniform distribution for the polar angle,

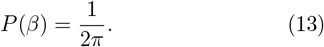

The autocorrelations of the polar variables vanish at nonzero lag, as shown in more detail in Appendix A,

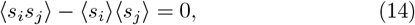

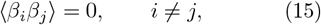

Thus, the polar representation naturally encodes the random walk in terms of i.i.d. random variables (*s*_*n*_, *β*_*n*_) sampled from a Rayleigh-distributed speed and a uniformly distributed orientation angle.

### C. Model 3: Random Walk in the comoving frame

The random walk can also be described in the comoving frame, where the relevant variables are the speed *s*_*n*_ and the turning angle *α*_*n*_. Since the orientation angles *β*_*n*_ are uniformly i.i.d. variables, the increments *α*_*n*_ defined by the linear Eq.(5) are also i.i.d. and uniformly distributed on the circle, see Appendix A for more details. These properties support a natural representation of the random walk in the comoving frame, defined by the pair (*s*_*n*_, *α*_*n*_) by sampling the speed i.i.d. as before from a Rayleigh distribution,

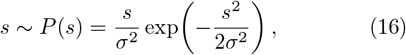

and the turning angle *α* i.i.d. from a uniform distribution on the circle, as originally suggested in Ref. [24],

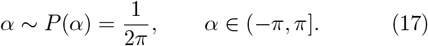

Here and in the following we map *α* onto the circle by applying a modulo 2*π* operation and restricting it to the interval (− *π, π*]. This choice corresponds to the minimal angular displacement and captures the tendency of an organism to change direction through the shortest possible rotation [61]. It also avoids a subtle issue of chirality in case we only admitted positive turning angles. The issue of wrapping onto the circle becomes more subtle for general stochastic processes, as we will discuss for the example of the OU process in Sec. IV. Thus, in the comoving frame, the random walk is governed again by two independent noise sources: Rayleigh noise in the speed and uniform white noise in the turning angle. A compact summary of the probability distributions and autocorrelation properties of all relevant variables in the three models (Cartesian, polar and comoving) is provided in Table I.

**TABLE I.** Statistical properties of different two-dimensional random walk variables in three different coordinate frames: Cartesian (Model 1), Polar (Model 2), and Comoving (Model 3).

| Model & Variables | Probability distributions | Autocorrelations |
| --- | --- | --- |
| **Model 1: Cartesian random walk** |  |  |
| Velocity components $(v_x, v_y)$ | $P(v_x, v_y) = \frac{1}{2\pi\sigma^2} \exp\left(-\frac{v_x^2 + v_y^2}{2\sigma^2}\right)$ | $\langle v_{x,i} v_{x,j} \rangle = \langle v_{y,i} v_{y,j} \rangle = 0$ for $i \neq j$ |
| **Model 2: Polar random walk** |  |  |
| Speed $s = \sqrt{v_x^2 + v_y^2}$ | $P(s) = \frac{s}{\sigma^2} \exp\left(-\frac{s^2}{2\sigma^2}\right), \quad s \geq 0$ | $\langle s_i s_j \rangle - \langle s_i \rangle \langle s_j \rangle = 0$ for $i \neq j$ |
| Orientation angle $\beta = \arctan(v_y, v_x)$ | $P(\beta) = \frac{1}{2\pi}, \quad \beta \in (0, 2\pi]$ | $\langle \beta_i \beta_j \rangle = 0$ for $i \neq j$ |
| **Model 3: Comoving random walk** |  |  |
| Speed $s$ | $P(s) = \frac{s}{\sigma^2} \exp\left(-\frac{s^2}{2\sigma^2}\right), \quad s \geq 0$ | $\langle s_i s_j \rangle - \langle s_i \rangle \langle s_j \rangle = 0$ for $i \neq j$ |
| Turning angle $\alpha = \beta_i - \beta_{i-1}$ | $P(\alpha) = \frac{1}{2\pi}, \quad \alpha \in (-\pi, \pi]$ | $\langle \alpha_i \alpha_j \rangle = 0$ for $i \neq j$ |

In summary, for a stochastic process defined in the Cartesian frame one needs to transform the corresponding probability distributions between the different frames, and in general to check for the correlation functions of the associated variables in the different frames (even though here for the Markovian random walk there is little to check). These analytical calculations are supported by numerical simulations. Figures 4,5 show that the probability densities of the Cartesian velocity components *v*_*x*_ and *v*_*y*_ are perfectly Gaussian for all three stochastic models, while the lower panels confirm that their autocorrelations vanish beyond zero lag, consistent with *δ*-correlated increments. Figure 6 compares the simulated probability densities of the speed with the analytical Rayleigh distribution Eq. (16) and demonstrates the absence of temporal correlations for all three models.

**FIG. 4.**
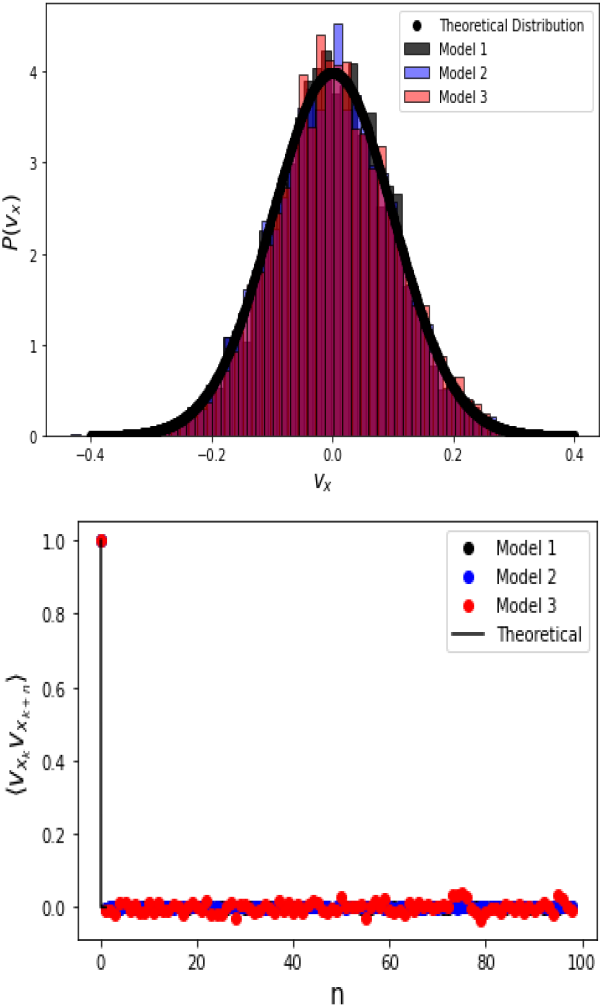
Top: Simulation results for the probability density of the velocity component *v*_*x*_ for the simple random walk formulated in three different frames, yielding three different random walk models, compared with the Gaussian distribution. Bottom: The autocorrelation function of the random walk velocities are all uncorrelated.

**FIG. 5.**
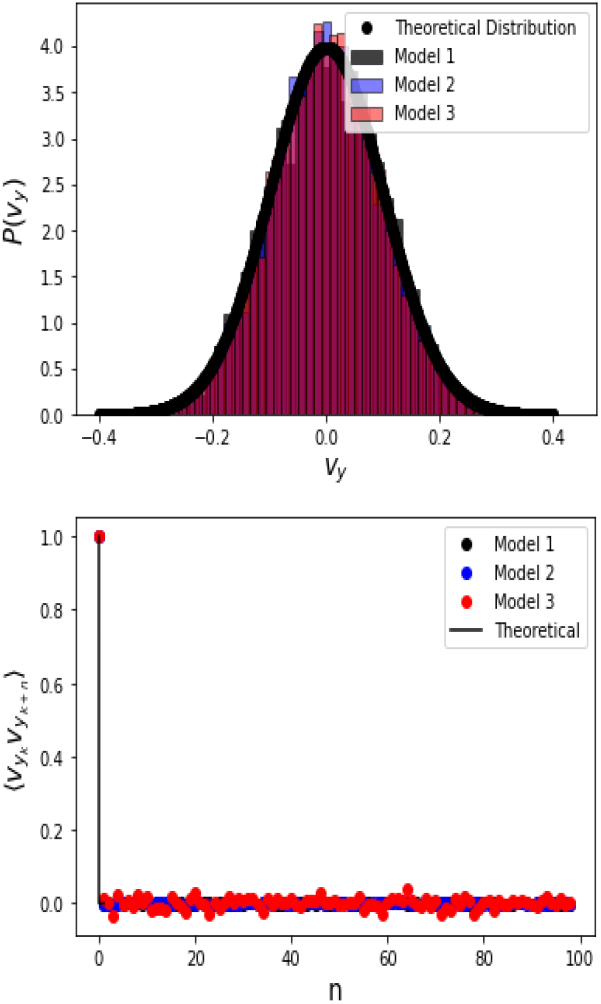
Same as Fig. 4 for the velocity component *v*_*y*_

**FIG. 6.**
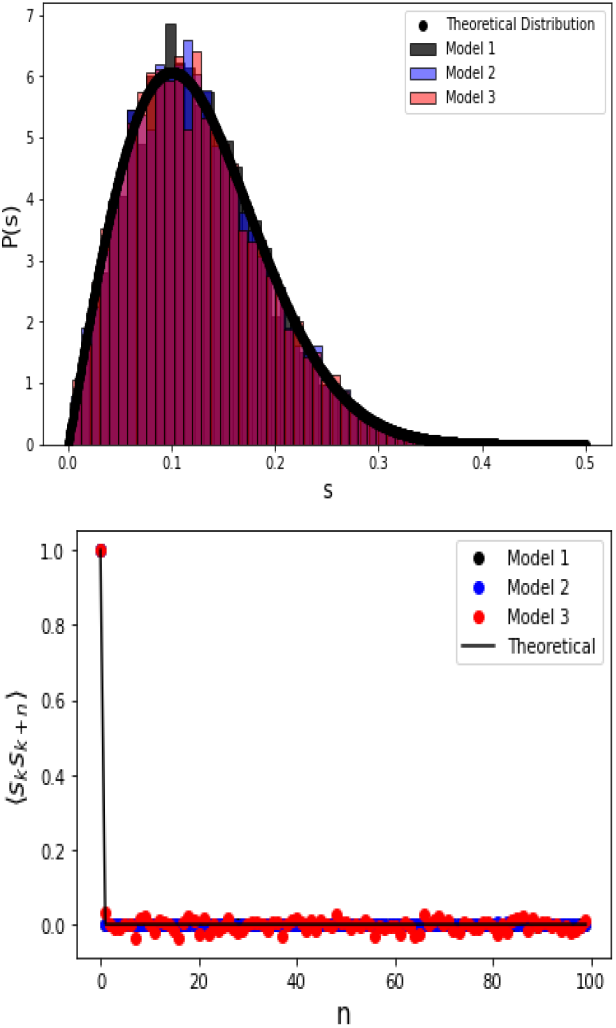
Top: Simulation results for the probability density of the speed *s* for the simple random walk in the three different frames compared with Eq.(16). Bottom: The autocorrelation function of the speed is again uncorrelated.

Similarly, Fig.7 shows the uniform distribution of the orientation angle *β* Eq. (13) and its lack of autocorrelation for all three models. Finally, Fig. 8 illustrates the distribution of turning angles *α*, which follows the uniform law in Eq. (17), and confirms its interpretation as a white noise process in angular space for all three models.

**FIG. 7.**
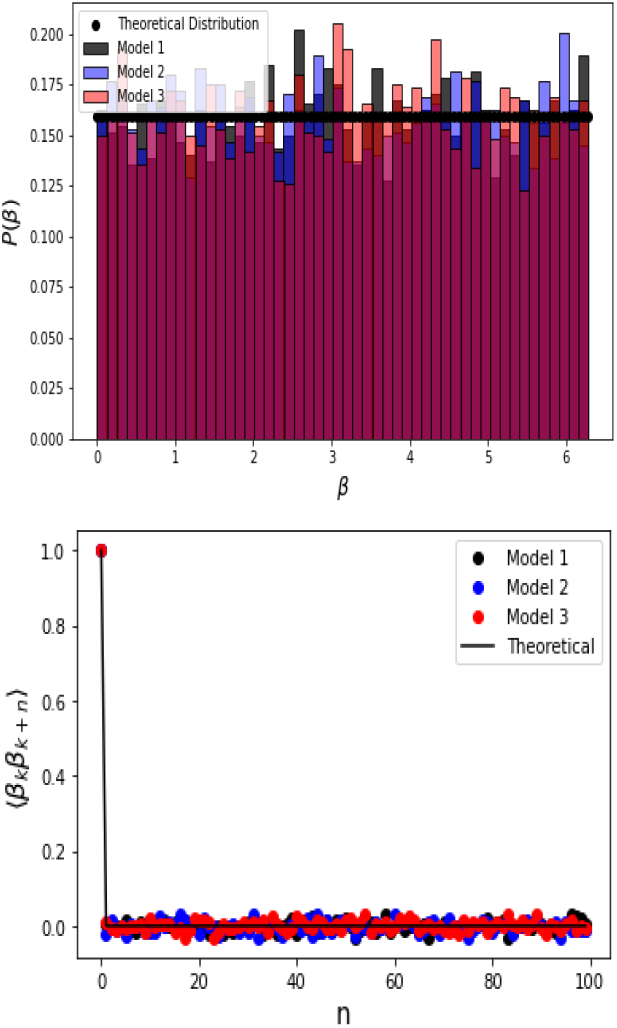
Top: Probability density of the orientation angle *β*_*n*_ for the simple random walk in the three different frames compared with Eq.(13). Below: The autocorrelation function is again uncorrelated.

**FIG. 8.**
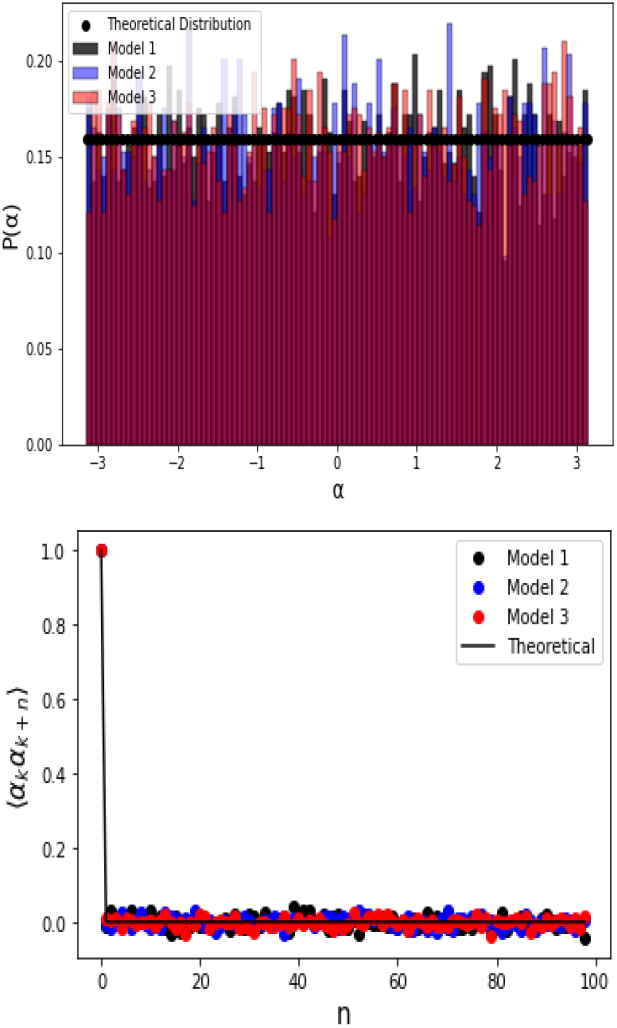
Top: Probability density of the turning angle *α*_*n*_ for the simple random walk in the three different frames compared with Eq.(17). Bottom: The autocorrelation function is again uncorrelated.

## IV. THREE MODELS FOR ORNSTEIN-UHLENBECK IN TWO DIMENSIONS

The conceptual framework of how to transform between the three different frames, illustrated in the previous section for the simple Markovian random walk, is now rolled out for the OU process yielding the velocities of random movements in a plane. Before elaborating on OU dynamics [38] in two dimensions, let us briefly recall its one-dimensional formulation and key statistical properties. The OU process is defined by the underdamped Langevin equation

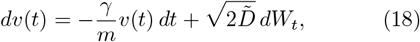

where *dW*_*t*_ is the increment of a Wiener process, *γ/m* is the relaxation rate, and the diffusion coefficient 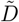 is related to the conventional diffusion coefficient *D* for the position via 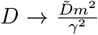 [62]. The solution of Eq. (18) is a Gaussian process with mean

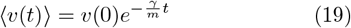

and variance

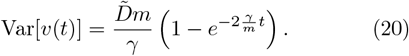

In the long-time limit, the process reaches a stationary state with Gaussian distribution

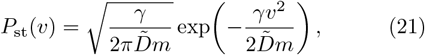

which has zero mean and variance *Dm/γ*. The velocity autocorrelation function of the stationary process is

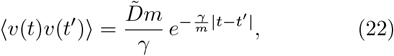

showing the characteristic exponential decay with correlation time *τ* = *m/γ*. The one dimensional OU process is therefore *Gaussian* (all finite-dimensional distributions are Gaussian), *stationary* (the distribution converges to the steady Gaussian Eq.(21)) and *Markovian* (the process has the Markov property, enabling Itô transformations to other coordinate systems [27]). These three properties make OU dynamics the prototypical model of a stochastic process with exponential memory. The two-dimensional OU process in Cartesian coordinates is then constructed as two independent copies of Eq. (18), one for each velocity component.

### A. Model 1: Cartesian Two-Dimensional OU Process

We first consider a two-dimensional OU-driven walk in the Cartesian frame, constructed from two independent one-dimensional OU processes for the velocities along the *x* and *y* axes. The velocity components **v** = (*v*_*x*_, *v*_*y*_) thus evolve according to the stochastic differential equation

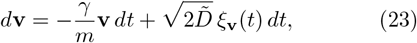

where 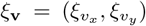 are independent Gaussian white noise terms.

The OU process exhibits exponentially correlated velocities with correlation function

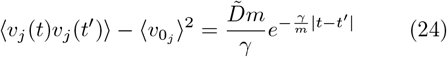

with *j* ∈ (*x, y*). The stochastic differential equation (23) is solved numerically in equilibrium using the Euler–Maruyama method [63] with parameters *D* = *γ* = *m* = 1 and noise terms *ξ*_**v**_ of zero mean and unit standard deviation. After integration, the corresponding polar frame variables are obtained via the polar transformation

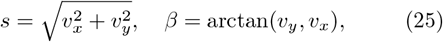

where *s* is the instantaneous speed and *β* the orientation angle.

We recall that the orientation angle *β*, extracted via the arctangent function, is defined modulo 2*π*, with values taken in the interval [0, 2*π*). Consequently, the turning angle *α* = Δ*β*, as a difference between these orientations, initially takes values on the full real line (−∞, ∞). Interestingly, this unwrapped turning angle (prior to any modulo operation) exhibits non-trivial temporal correlations. Wrapping *α* onto the circle, as by definition needed for comoving frame dynamics, eliminates these correlations, thus effectively Markovianising the turning angle dynamics. This represents a loss of information in the comoving frame about the history of previous rotations with respect to the Cartesian frame.

### B. Model 2: Polar OU Process Driven by Itô Equations

Defining stochastic equations in the polar frame introduces an Itô–Stratonovich ambiguity, as the transformation from Cartesian velocities to polar coordinates generates multiplicative noise for the speed *s* and orientation *β*. Using Itô calculus [27], Eq. (23) can be transformed into two coupled stochastic differential equations for the speed *s* and orientation *β* (see Appendix B), as similarly derived in Ref. [27] for an electric field, and for the Stratonovich approach in Ref. [47]:

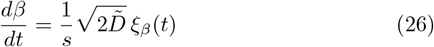

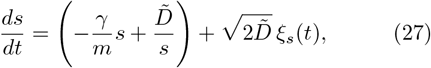

where the noise terms are defined as

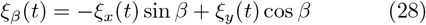

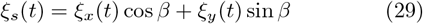

with *ξ*_*x*_ and *ξ*_*y*_ being the independent Gaussian white noises in the Cartesian frame. It can be shown that *ξ*_*β*_ and *ξ*_*s*_ are also Gaussian white noises [27]. The corresponding Fokker–Planck equation [64] for the joint probability density function *P*(*s, β, t*) in the Itô interpretation is

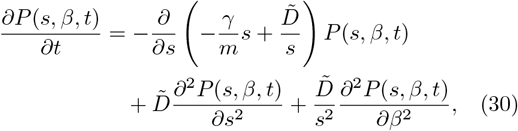

which admits the stationary solution

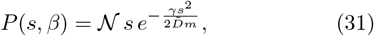

where 𝒩 represents the uniform distribution of *β* and the speed *s* follows a Rayleigh distribution. For simulations, Eqs. (26) and (27) are integrated using the Euler–Maruyama method [63], ensuring that the standard deviations of *ξ*_*s*_ and *ξ*_*β*_ match those of the Cartesian noise in Model 1. A key consideration arises from Eq. (27): Numerically, the Gaussian noise in *ξ*_*s*_ can occasionally produce negative values for the speed. To maintain positive speeds, any negative speed values encounter in the simulation, are replaced by a small threshold value of 10^−2^. Note that analytically negative speeds Eq. (27) appear to be eliminated by the Itô 1*/s* flux term, although we have no proof of the positivity of the speed in this equation.

The autocorrelation function of the speed can be calculated (see Appendix C) as

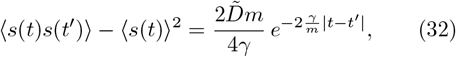

which is analogous to the velocity correlations in the Cartesian frame Eq. (24) but decays at twice the rate, 2*γ/m*, reflecting the distinct dynamics in polar coordinates.

Deriving a closed-form expression for the autocorrelation of *β* is more involved due to its inverse dependence on *s*, which introduces a nontrivial coupling between speed and orientation. Consequently, instead we consider the autocorrelation of cos *β*, which can be approximated as discussed in Appendix D to obtain

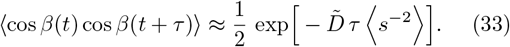

### C. Model 3: OU Process in the Comoving Frame

To achieve a self-consistent description of the two-dimensional OU process in the comoving frame, it is necessary to formulate an equation for the turning angle *α*. This can be accomplished by using Eq. (5) and computing the corresponding distribution of *α* from the righthand side of Eq. (26), restricting the turning angle to the interval [− *π, π*] with the modulo 2*π* operation. The resulting turning angle probability distribution is given accordingly to (see Appendix F)

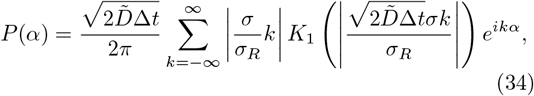

where Δ*t* is the discretisation time for the originally time-continuous Eq. (5), and *σ, σ*_*R*_ are the standard deviations of the Gaussian white noise *ξ*_*β*_ (from Eq. (26)) and the exponentially correlated speed *s*(*t*) (from Eq. (31)), respectively. Here, *K*_1_(*z*) denotes the modified Bessel function of the second kind. While *P*(*α*) resembles a wrapped Gaussian or von Mises distribution as shown in Fig. 9, its precise functional form defines, to our knowledge, a new distribution for the turning angle.

**FIG. 9.**
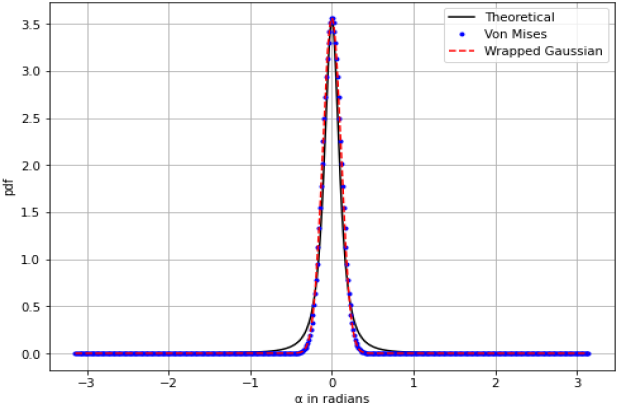
Theoretical probability distribution for the turning angle *α*, Eq. (34), compared with a wrapped Gaussian and a von Mises distribution. One can see clear deviations in the tails.

In a steady state, the speed *s* is distributed according to a Rayleigh distribution with exponentially decaying autocorrelations as in Eqs. (31),(32). This leads to a self-consistent formulation of a Cartesian OU process represented in the comoving frame as

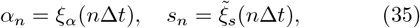

where *ξ*_*α*_ is white noise with the functional form given by *P*(*α*), and 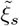 is exponentially correlated Rayleigh noise. Alternatively, the speed can be generated using Eq. (27) with Gaussian white noise. What we mean with a self-consistent description of the OU process in the comoving frame is that the above two equations are formulated solely in terms of speed and turning angle, without explicitly involving any other coordinates. This is not the case in the polar representation, which still somewhat relies on the Cartesian frame. Note that, exactly in this sense, here we go beyond previous formulations of active particle models, such as in Refs. [47, 48, 53, 54], where the active part is expressed in terms of polar coordinates but not defined self-consistently by using the turning angle as a comoving frame coordinate, as above. In principle, for a walker, or an organism, knowledge of a sequence of speed and turning angle is fully sufficient to self-generate movement, which is the idea underlying Pearson’s original random walk and the CRW of ME. However, to our knowledge, generalised processes of the form of Eq. (35) have not yet been considered for modelling organismic movements.

For simulations, the variables in the comoving frame can be generated directly from their respective distributions, i.e., the Markovian turning angle *α* is sampled i.i.d. from *P*(*α*) Eq.(34). However, to transform back to the polar or Cartesian frames by using Eq.(6) for the orientation angle *β*, the correct time discretisation taking the diffusive stochastic scaling into account has to be considered [27]. For this we introduce an additional helper variable, the angular velocity *ω* = *dβ/dt*, thus writing 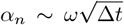, where *ω* is sampled from the probability distribution for the angular velocity associated with the turning angle distribution, see Eq. (F13) in Appendix F. Simultaneously, the speed *s* is sampled from exponentially correlated Rayleigh noise. This procedure maintains the correct temporal correlations and steady-state statistics of the Cartesian OU process. Correlations for *s* can be introduced numerically using the Cholesky decomposition method [65], generating two Gaussian variables with exponential correlations as in Eq. (24). According to [66], if *X* and *Y* are independent Gaussian variables, then 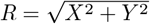 follows a Rayleigh distribution with standard deviation *σ*_*R*_. In this manner, exponentially correlated Rayleigh noise is obtained. Along these lines, once the comoving variables (*α, s*) are generated, the transformations in Eq. (5), modulo using *ω*, are applied to recover the Cartesian velocities.

This description completes our definition of the three models of the OU process in the three different coordinate frames. We now compare results from simulating these three models with each other, and with analytical results, to demonstrate that all these different formulations reproduce the same statistical properties in the different frames. The simulation results presented in Figs. 10, 11, 12, and 13 show the probability distributions and associated autocorrelation functions for each key variable: speed *s*, turning angle *α*, orientation angle *β*, angular velocity *ω*, and velocities *v*_*x*_, *v*_*y*_, as summarized in Table II. The figures demonstrate strong agreement between all three models. The probability distributions for each variable exhibit very similar shapes and peak locations, indicating that the underlying statistical properties of the simulated trajectories are well preserved. Likewise, the autocorrelation functions decay at comparable rates, demonstrating that temporal correlations and memory effects are consistently captured across the different model implementations. For instance, Fig.10 presents the speed distributions and autocorrelations in the comoving frame, which closely match the theoretical Rayleigh distribution and the exponential decay predicted by Eq.(32). Figure 11 shows the turning angle distributions compared with Eq. (34) and their approximately *δ*-like autocorrelations. The orientation angle *β* distributions and correlations are displayed in Fig.12. We remark again that obtaining an analytical expression for the autocorrelation of *β* looks impossible due to its dependence on the speed in Eq.(26). Instead, we therefore consider our analytical approximation for the autocorrelation of cos *β*. A small mismatch is observed in these results, which can be attributed to two main factors: (i) when sampling the random variables in the comoving frame from Eq. (34), the infinite discrete Fourier series must be truncated at a finite value *N* due to the computational cost of simulating large terms. This truncation directly affects the accuracy of the sampled variables. (ii) The choice of the simulation step size Δ*t* influences the recovery of continuous trajectories, i.e., smaller steps improve the accuracy but increase computational expense. Finally, the Cartesian velocity components *v*_*x*_ and *v*_*y*_, illustrated in Fig.13, confirm Gaussian behavior with temporal correlations consistent with theoretical predictions, further validating the equivalence of the constructed three models.

**TABLE II.**
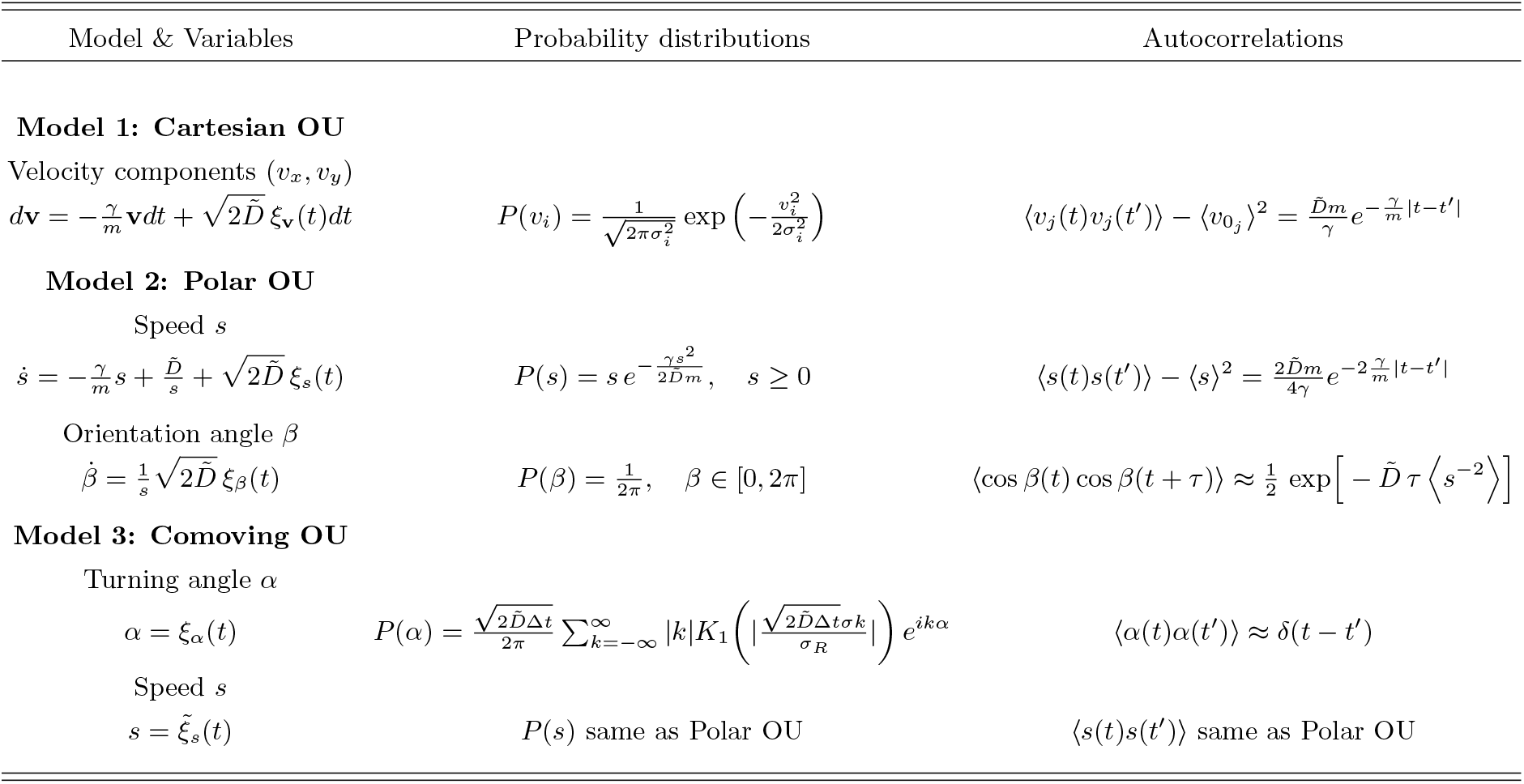
Statistical properties of different two-dimensional OU process variables in three different coordinate frames: Cartesian (Model 1), Polar (Model 2), and Comoving (Model 3).

**FIG. 10.**
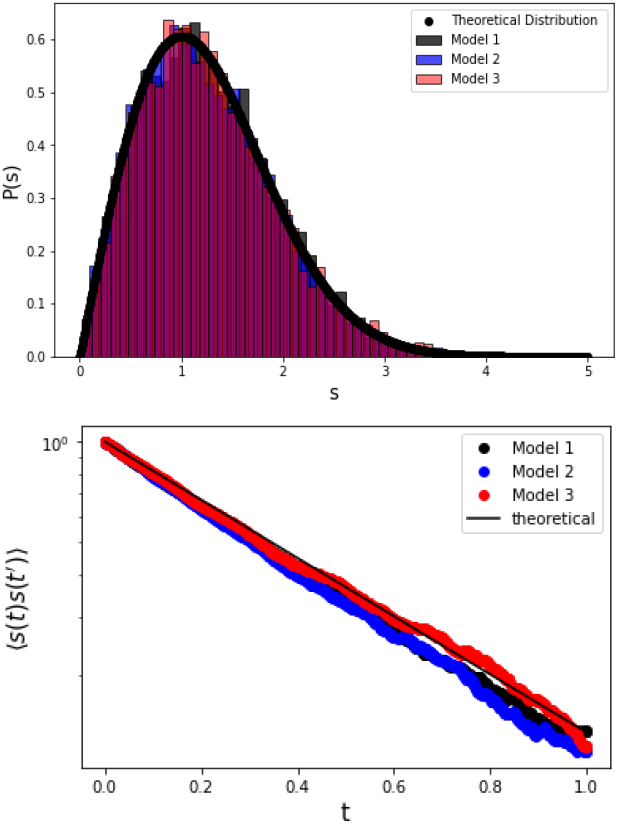
Top: Probability distributions for the speed *s* in the three different models of an OU process in two dimensions, compared with Eq.(31) (lines). Bottom: Corresponding speed autocorrelations compared with Eq.(32) for the three different models. Here and in the following *t* represents the total time length of the process.

**FIG. 11.**
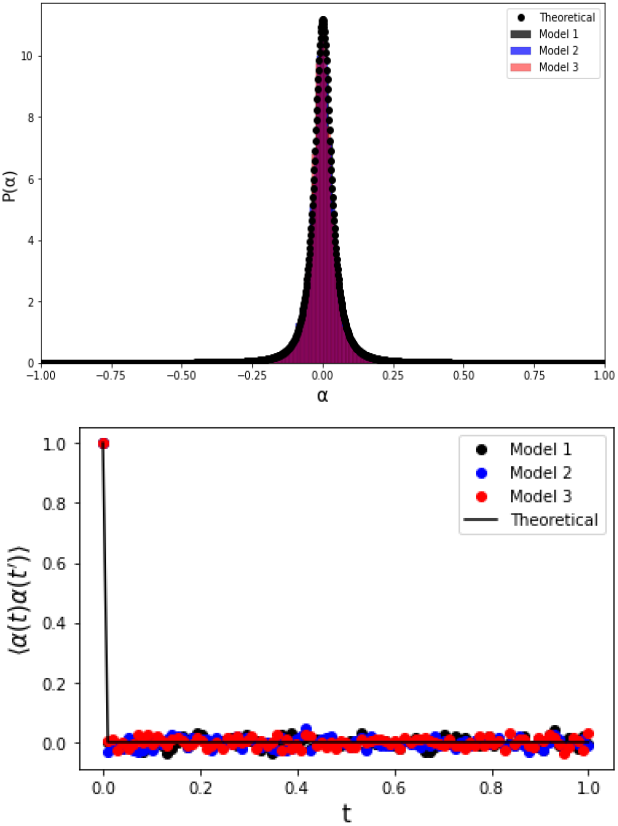
Top: Probability distributions for the turning angle *α* in the three different models of an OU process in two dimensions, compared with Eq.(34) (lines), the original distribution is defined in the interval [− *π, π*), but a focus is made in the interval [− 1, 1] for visual purposes. Bottom: Corresponding autocorrelations for the three different models, compared with a *δ* function.

**FIG. 12.**
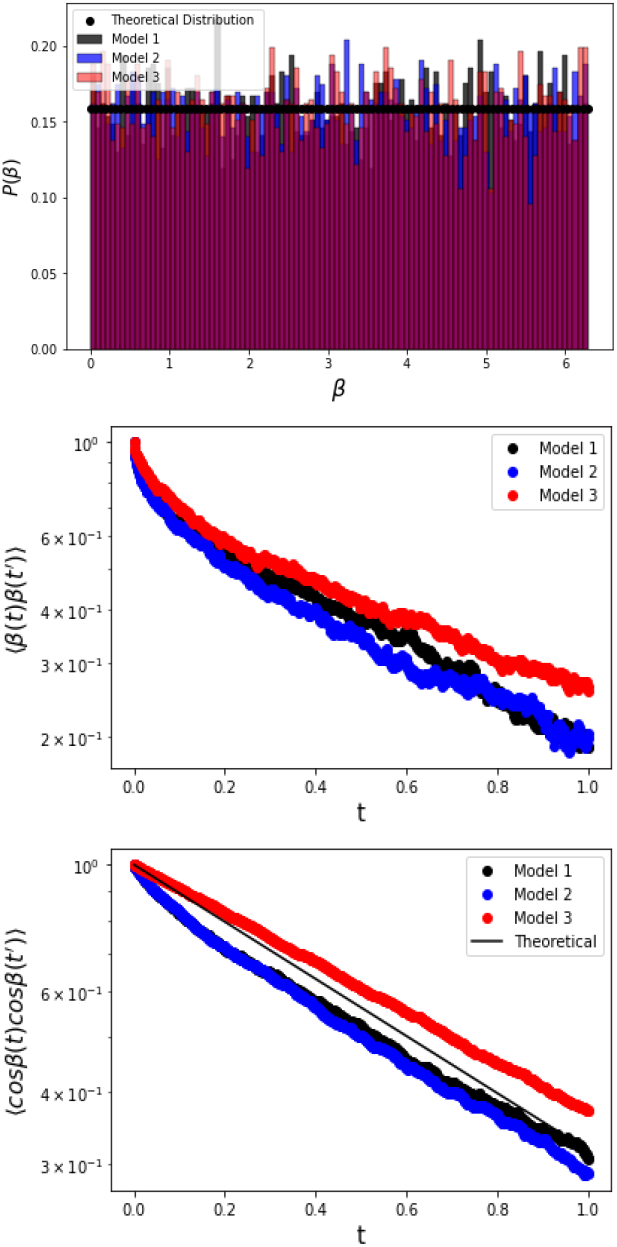
Top: Probability distributions for the orientation angle *β* in the three models, compared to a uniform distribution. Middle: Corresponding orientation autocorrelations for the three models. Bottom: Corresponding autocorrelations for the cosine of the orientation, including the analytical approximation Eq. (D4) (line).

**FIG. 13.**
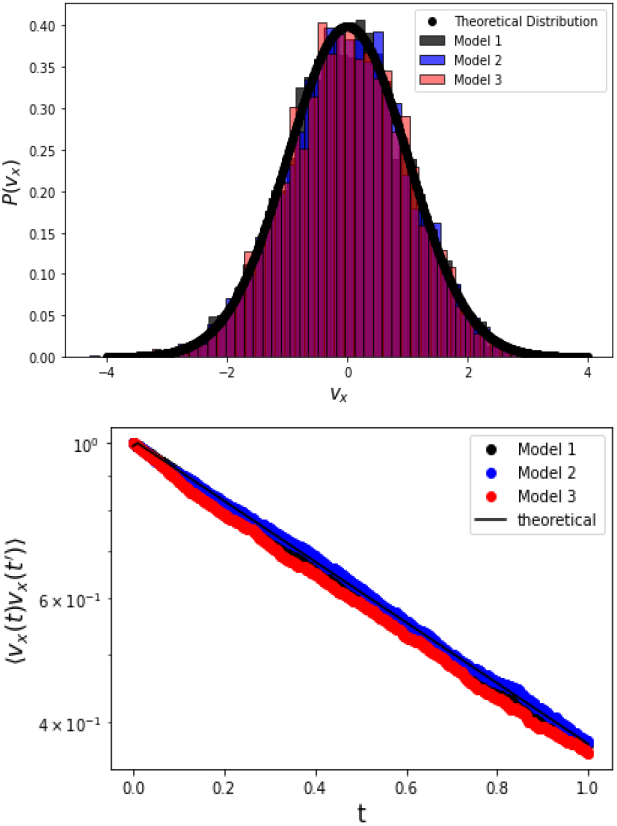
Top: Probability distributions for the velocity *v*_*x*_ in the three models compared with the corresponding theoretical expression for a Gaussian distribution (line). Bottom: Correspondind velocity autocorrelations compared with the theoretical expression (line) in Eq.(22).

Overall, our comparative analysis confirms that all three models consistently numerically reproduce the statistical properties of the two-dimensional Cartesian OU process across all frames. As far as we can tell, our analytical transformations between Cartesian, polar, and comoving frames are exact, which is confirmed by the overall excellent agreement of both probability distributions and autocorrelation functions.

## V. SUMMARY AND CONCLUSIONS

In summary, what we called the comoving frame is nothing else than the coordinate frame suggested by Ross and Pearson to formulate a two-dimensional random walk for modelling organismic movements [21–23]. From these early origins comoving coordinates propagated into ME in the form of the widely used stochastic CRW model [2, 30]. Turning angle and step length distribution functions are in turn much extracted from experimental data for moving organisms [50–52]. This theoretical framework is at variance with stochastic models in active matter theory, which are all formulated in the Cartesian coordinate frame, partially by using polar coordinates [47, 48, 53, 54]. However, as we pointed out in this article, the polar frame should not be confused with the comoving frame. Defining stochastic dynamics in the latter frame involves a non-trivial additional step beyond polar coordinates, which is deriving a stochastic equation for the turning angle dynamics, in addition to the speed dynamics. The important advantage of the comoving frame is that it allows to formulate a set of equations which more closely mimicks actual biological movements by describing the fluctuations generating them in terms of equations that are self-consistent in this frame. This approach matches to the biophysical reasoning that self-propelled biological movements are generated internally by an organism itself, and not by any external noise in a fixed coordinate frame. Such a fixed Cartesian frame should only be used to model the impact of external environmental conditions on a moving organism. If a given stochastic process can be transformed exactly between the Cartesian and the comoving frame, of course it does not matter in which frame one expresses the given dynamics; however, as we will argue further below, exact transformations might not always be possible.

In our article, we have first defined the relevant three different frames, with Cartesian and polar being trivial, as explained in textbooks, but by identifying the comoving frame on this basis. The essential idea of transforming stochastic processes between these three different frames was first outlined for the simple random walk as an example, and on this basis worked out in detail for the OU process. While spherical and polar transformations of stochastic dynamics and associated representations of OU probability distributions are well known [47, 53, 67], to our knowledge the autocorrelation functions for OU speed and orientation angle have not been obtained before, which constitute first new results. A more important finding is our, as far as we can tell, novel turning angle distribution calculated exactly analytically for the OU process, which does not seem to match to other famous circular distributions like wrapped Gaussian or van Mises distributions.

Our most important result, however, is the self-consistent formulation of the Cartesian OU process in the comoving frame. We argue that our derivation is exact, as is in line with our numerical results. Interestingly, our comoving OU model generates a unimodal turning angle distribution, as is characteristic for the CRW of ME. However, on top of this it features an exponentially decaying autocorrelation function for the speed. To our knowledge, such a generalised stochastic model defined in the comoving frame has not yet been studied in ME, nor tested for organismic movements, even though the importance of correlation functions to understand organismic movements has been emphasized before [68].

Importantly, the comoving OU equations that we have derived form a special case of the generalised Langevin dynamics in the comoving frame stipulated *ad hoc* in Ref. [3], reading

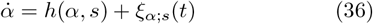

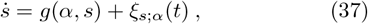

where the forces on the right hand sides are split into coupled drift terms and general noises. Considering an overdamped version of Eq. (36) with a drift term that is linear in *α* reproduces the form derived for OU of Eq. (35) for the turning angle [3]. The associated underdamped equation for the speed clearly yields a special case of Eq. (37). Our results for the comoving OU process thus demonstrate that the theoretical framework heuristically provided by Eqs.(36),(37) can be established, at least for the special case of the OU process, from first principles. To our knowledge, this is the first time that a comoving OU process has been derived. Hence, our second main results is to have established, along the lines of the OU process, a general theoretical framework that clearly defines, and disentangles, the different coordinate frames for generally transforming stochastic processes from the Cartesian into the comoving frame. Our approach can thus be used to systematically construct more general stochastic movement models than CRWs in the comoving frame.

Especially, along these lines it would be interesting to merge the approach put forward by active matter theory in terms of active particle models with CRW models of ME [1]. Generalised active particle models combining different types of stochastic dynamics have already been considered recently [69]. In forthcoming work we will show, by exploiting the conceptual framework developed in this article, how to transform three generic active particle models into the comoving frame. This will pave the way to develop more general active particle models, and to make these ideas attractive to applications in ME.

However, our present conjecture is that exact transformations between the three different frames are only possible if the Cartesian stochastic dynamics is Markovian, as for simple random walks and the considered OU process. There is indeed a non-trivial Markovianisation taking place viz. a loss of memory in the turning angle dynamics when being wrapped onto the circle. It is indeed well known already that there exist three different semi-Markovian Lévy walk models, two defined in the Cartesian frame and one in the comoving frame, which cannot be transformed into each other [70]. This result provides first evidence that it may in general not be possible to transform non-Markovian processes defined in the Cartesian frame exactly into the comoving frame. Indeed, for fractional Brownian motion a comoving representation is, to our knowledge, not known [56]. However, there is a need to formulate representations of more advanced stochastic processes in the comoving frame for implementing self-consistent stochastic navigation in robots and drones [71].

Interestingly, if exact transformations of stochastic processes with memory between these different frames are not possible, one might conclude that active fluctuations generating self-propelled organismic movements should *per se* not be defined in the Cartesian or polar frames but exclusively self-consistently in the comoving frame. This would, in turn, necessitate to develop a novel theory of stochastic processes in the comoving frame, and to explore how such processes then look like in the Cartesian frame. We finally remark that the relevance of distinguishing between these different frames of reference is already well-known to biologists studying organismic movements. They call a world-centered frame, corresponding to the Cartesian one, *allocentric* while the body-centered, comoving one is known as *egocentric* [72]. In the former, animals navigate according to fixed external landmarks while in the latter, they integrate their path coordinates internally in the brain [72, 73]. These two frames are, in turn, represented in the brain by different types of neurons, grid and place cells, respectively [74]. There are thus plenty of reason to further explore the theory of stochastic process in the comoving frame.

## ACKNOWLEDGMENTS

NLA wish to acknowledge to “Fundacion Politecnico” and the School of Mathematical Sciences from Queen Mary University of London for the financial support of this research. NLA and RK thank Alessia Gentili, Giorgio Volpe, Adrian Baule and Wolfram Just for very helpful discussions.

## Appendix A Autocorrelations in Speed and Orientation Random Walk

We consider a two–dimensional random walk defined by

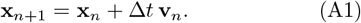

Since the velocities **v**_*n*_ are i.i.d. for different time indices, and since (*s*_*n*_, *β*_*n*_) are deterministic functions of **v**_*n*_, it follows that

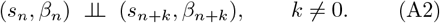

### 1. Autocorrelation of the speed

The autocovariance of the speed is defined as

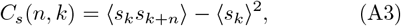

where the angular brackets denote an ensemble average. For *n* ≠ 0, temporal independence (A2) implies

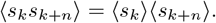

By stationarity, ⟨*s*_*k*_⟩ = ⟨*s*_*k*+*n*_⟩ = ⟨*s*⟩, and therefore

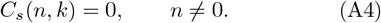

### 2. Autocorrelation of the orientation angle

The autocovariance of the orientation angle is defined as

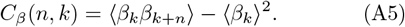

For *n* ≠ 0, Eq. (A2) yields

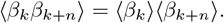

Since *β* is uniformly distributed in [0, 2*π*),

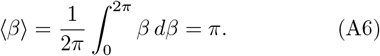

Consequently,

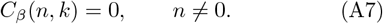

#### 3. Autocorrelation of the turning angle

We define the turning angle as the difference between two successive orientation angles,

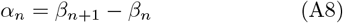

with *α*_*n*_ ∈ (−*π, π*]. Because *β*_*n*+1_ and *β*_*n*_ are independent and uniformly distributed, the turning angle *α*_*n*_ is itself uniformly distributed on (− *π, π*]. Moreover, for *n* ≠ 0, the pairs (*β*_*k*+1_, *β*_*k*_) and (*β*_*n*+*k*+1_, *β*_*n*+*k*_) are independent. Therefore,

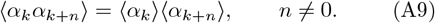

The autocovariance of the turning angle,

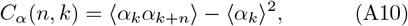

thus vanishes for all nonzero lags,

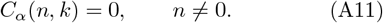

## Appendix B Itô transformation

We start with the two-dimensional stochastic differential equation for the velocity **v** = (*v*_*x*_, *v*_*y*_):

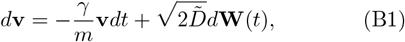

where *d***W**(*t*) = (*dW*_*x*_, *dW*_*y*_) is a vector of independent Wiener processes.

We define the polar coordinates:

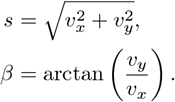

We apply Itô’s lemma to find the stochastic differential equations for *s* and *β*. Let 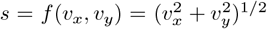.

The partial derivatives are:

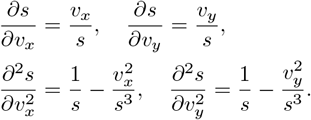

The drift and diffusion coefficients for *v*_*x*_ and *v*_*y*_ are:

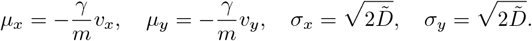

By Itô’s lemma:

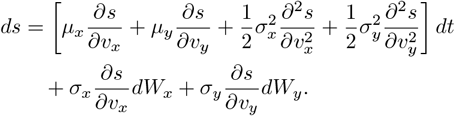

Substituting the derivatives:

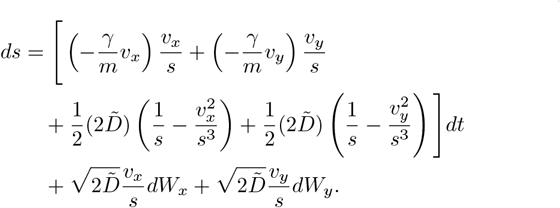

Simplifying the drift term:

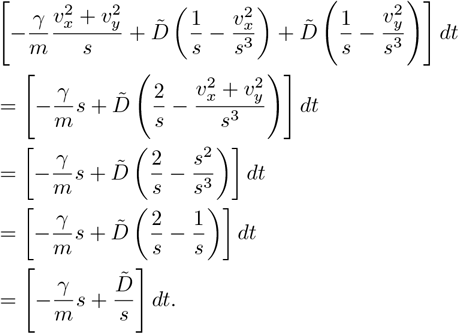

The diffusion term is:

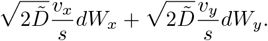

This can be written as 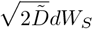, where 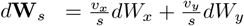 is a Wiener process.

Thus, the stochastic differential equation for *S* is:

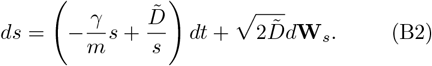

Let *β* = *f*(*v*_*x*_, *v*_*y*_) = arctan 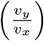. The partial derivatives are:

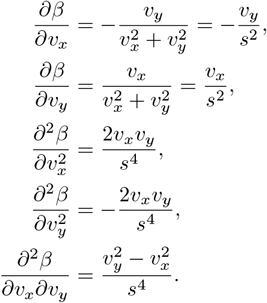

By Itô’s lemma:

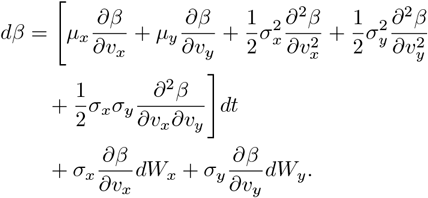

Since 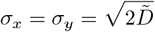 and the noises are independent, the cross term is zero. Substituting:

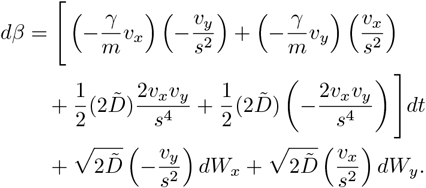

The drift terms cancel:

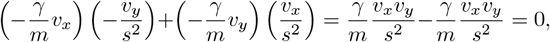

and

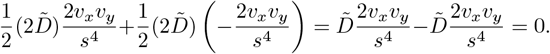

Thus, the drift is zero.

The diffusion term is:

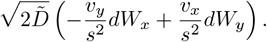

This can be written as 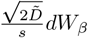, where 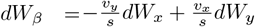 is a Wiener process.

Thus, the stochastic differential equation for *β* is:

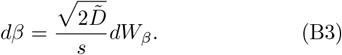

In summary, the transformed stochastic differential equations in polar coordinates are:

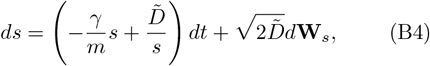

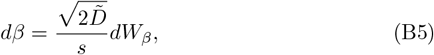

where *dW*_*S*_ and *dW*_*β*_ are independent Wiener processes.

## Appendix C Autocorrelation Function for the Speed in th OU

To obtain the mean value of the speed, we examine the steady-state behavior of Eq.(27). In this limit, the left-hand side becomes zero:

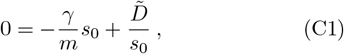

where *s*_0_ represents the mean speed in the steady-state. Solving for *s*_0_ we find:

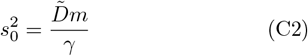

which implies that 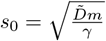.

Secondly, to calculate the autocorrelations for *s*, we introduce a linear expansion around *s*_0_ for *s*(*t*):

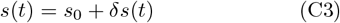

where *δs*(*t*) represents the fluctuations around the steady-state value *s*_0_. Substituting this into Eq.(27):

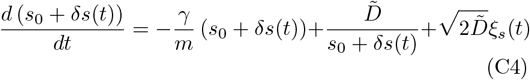

From this, we observe that the term 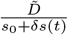 can be rewritten as:

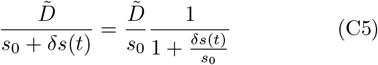

Here, we can expand the factor 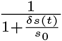 in a Taylor series for small values of 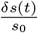:

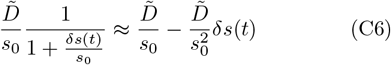

Thus, Eq.(C4) becomes:

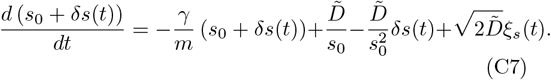

Applying the linearity of the differential operator on the left side and substituting 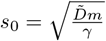, several terms cancel out:

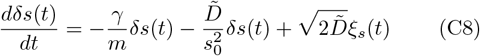

Simplifying, we get:

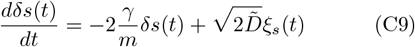

We notice that Eq.(C9) has the same form as an OU process. Hence, its autocorrelation can be calculated similarly as:

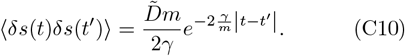

Now, for our original variable *s*(*t*), the autocorrelation is:

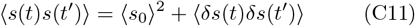

Finally, we can compute the autocorrelation function:

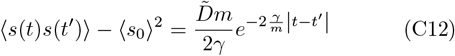

## Appendix D Autocorrelation of cos *β* in OU

Consider the orientation angle *β*(*t*) evolving according to the polar OU process:

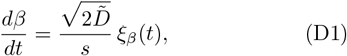

where *ξ*_*β*_(*t*) is Gaussian white noise with zero mean and unit variance. For a discrete time increment Δ*t*, the angular increment can be approximated as

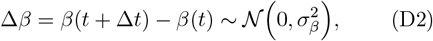

with variance

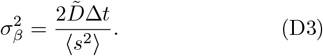

With the assumption of Δ*β*(*t*) as Gaussian distributed, the autocorrelation of cos *β* is then

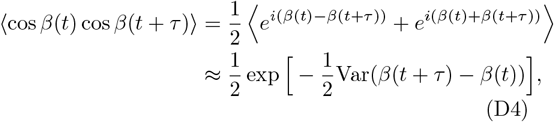

Combining with (D3) gives

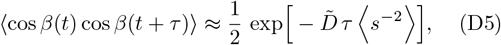

where ⟨*s*^−2^ ⟩ can be approximated by numerical fitting to the simulations of the models. This expression shows that the autocorrelation of cos *β* decays exponentially, with a rate determined by both the OU relaxation time and the variance of the angular increments 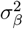.

## Appendix E Autocorrelation of *ω*(*t*) and *α* in OU

The autocorrelation of *ω*(*t*) is defined as:

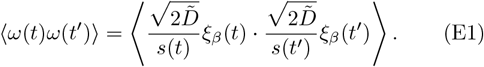

Using the property of white noise ⟨ *ξ*_*β*_(*t*)*ξ*_*β*_(*t*^*′*^) ⟩ = *δ*(*t* − *t*^*′*^), we simplify the expression:

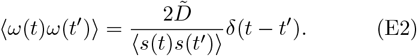

which shows that the angular velocity is simply a delta correlated variable. Accordingly, for the turning angle, whi<u>ch</u> is directly related to the angular velocity via 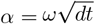, we obtain that the autocorrelations are

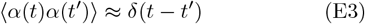

## Appendix F Calculation of the Probability Distribution for the Turning Angle in the OU

According to the Itô equation (26), the instantaneous angular velocity 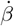 is given by the term 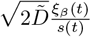. To calculate the probability distribution of the turning angle *α*, we note that 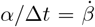. Therefore, the scale parameter of the resulting probability density function (PDF) will be modified by the time step Δ*t*. The steps to find such a PDF are as follows:

First, we perform a change of variables for the term 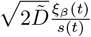

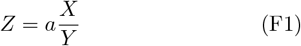

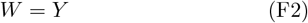

where *X* represents a Gaussian white noise with standard deviation *σ, Y* represents an exponentially correlated Rayleigh distribution with scale parameter *σ*_*R*_, and *a* is a constant. The joint distribution *P*(*z, w*) can be found using the Jacobian of the transformation:

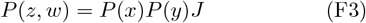

The Jacobian *J* is calculated as:

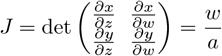

Now, we can write the joint distribution as:

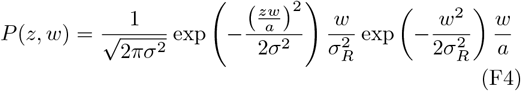

which can be simplified to:

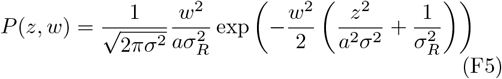

Since we are interested in the marginal distribution for *z*, we integrate the joint distribution with respect to *w* from 0 to ∞:

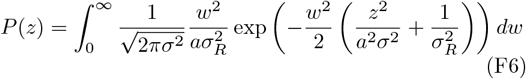

Upon integration, we find that:

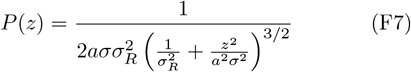

Simplifying this expression yields:

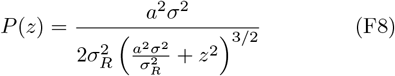

This is the PDF for the variable on the real line in the comoving frame. However, for describing particle turning angles, a representation within the range [− *π, π*] is more natural. To achieve this, we wrap our PDF to the circle by applying the Poisson Summation Formula:

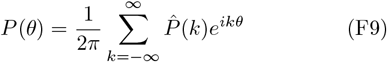

where 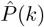 is the Fourier transform (FT) of *P*_*Z*_(*z*) from Eq.(F8). We calculate the FT:

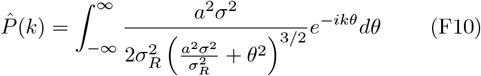

This integral evaluates to:

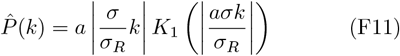

where *K*_1_(*z*) is the modified Bessel function of the second kind of order 1. Now, substituting this into the Poisson formula:

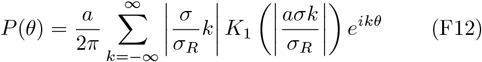

This represents the PDF for the ratio of a Gaussian distribution and a Rayleigh distribution in the comoving frame, within the range [−*π, π*].

Substituting 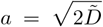 to find the angular velocity PDF:

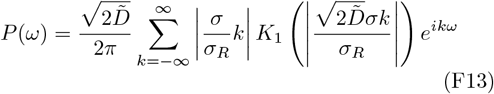

Here, the variable *ω* is in the interval [− *π, π*] with units of radians per second (rad/s), as this is an angular velocity.

In the case of the PDF of the turning angle *α*, we need to substitute 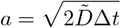, obtaining:

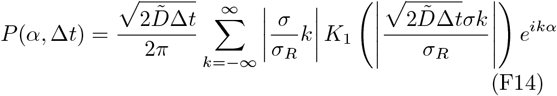

Here, we observe a dependence on the time step Δ*t* chosen for the simulations, as this refers to the turning angle, which will have a different distribution depending on how frequently the dynamics are sampled. For this case, the interval is between [−*π, π*] with units of radians.

